# Pangenomic analysis of *Helcococcus ovis* reveals widespread tetracycline resistance and a novel bacterial species, *Helcococcus bovis*

**DOI:** 10.1101/2024.05.20.594939

**Authors:** Federico Cunha, Yuting Zhai, Segundo Casaro, Kristi L. Jones, Modesto Hernandez, Rafael S. Bisinotto, Subhashinie Kariyawasam, Mary B. Brown, Ashley Phillips, Kwangcheol C. Jeong, Klibs N. Galvão

**Affiliations:** Department of Large Animal Clinical Sciences, University of Florida College of Veterinary Medicine, Gainesville, Florida; Department of Animal Sciences, University of Florida College of Agriculture and Life Sciences, Gainesville, Florida; Emerging Pathogens Institute, University of Florida, Gainesville, Florida; D. H. Barron Reproductive and Perinatal Biology Research Program, University of Florida, Gainesville Florida; Department of Comparative, Diagnostic, and Population Medicine, University of Florida College of Veterinary Medicine, Gainesville, Florida; Department of Infectious Diseases and Immunology, University of Florida College of Veterinary Medicine, Gainesville, Florida; Athens Veterinary Diagnostic Laboratory, College of Veterinary Medicine, The University of Georgia, United States

## Abstract

*Helcococcus ovis* (*H. ovis*) is an opportunistic bacterial pathogen of a wide range of animal hosts including domestic ruminants, swine, avians, and humans. In this study, we sequenced the genomes of 35 *Helcococcus sp.* clinical isolates from the uterus of dairy cows and explored their antimicrobial resistance and biochemical phenotypes. Phylogenetic and average nucleotide identity analyses placed four *Helcococcus* isolates within a cryptic clade-representing an undescribed species, for which we propose the name *Helcococcus bovis* sp. nov. We applied whole genome comparative analyses to explore the pangenome, resistome, virulome, and taxonomic diversity of the remaining 31 *H. ovis* isolates*. H. ovis* was more often isolated from cows with metritis, however, there was no associations between *H. ovis* gene clusters and uterine infection. The phylogenetic distribution of high-virulence determinants of *H. ovis* is consistent with convergent gene loss in the species. The majority of *H. ovis* strains (30/31) contain mobile tetracycline resistance genes, leading to higher minimum inhibitory concentrations of tetracyclines in vitro. In summary, this study showed that the presence of *H. ovis* is associated with uterine infection in dairy cows, that mobile genetic element-mediated tetracycline resistance is widespread in *H. ovis*, and that there is evidence of co-occurring virulence factors across clades suggesting convergent gene loss in the species. Finally, we introduced a novel *Helcococcus* species closely related to *H. ovis*, called *H. bovis* sp. nov.

**Highlights:**

- The presence of *Helcococcus ovis* is associated with uterine infection in dairy cows
- Mobile genetic element-mediated tetracycline resistance is widespread in *H. ovis*
- Co-occurring virulence factors across clades suggest convergent gene loss in the species
- *Helcococcus bovis* is a novel species closely related to *Helcococcus ovis*

## Introduction

*Helcococcus ovis (H. ovis)* belongs to a clinically important genus of bacteria populated by five other described species: *Helcococcus kunzii*, *Helcococcus massiliensis*, *Helcococcus sueciensis*, *Helcococcus seattlensis*, and *Helcococcus pyogenes*, all of which are opportunistic pathogens of humans (1–4). Unlike the remaining members of the genus, *H. ovis* is most often found as a co-infecting pathogen in mixed infections of farm animals, such as metritis (5), mastitis (6), and pneumonia (7). Due to these characteristics, its ability to independently cause disease had not been documented until recently (7–10). *H. ovis* is capable of independently causing bovine valvular endocarditis (11), pneumonia, and bursitis (12) in clinical infections, and mastitis in an experimentally infected mouse (*Mus musculus*) model (6). The only confirmed human infection by *H. ovis* originates from a 2018 case study of an artificial eye infection caused by the then-named Tongji strain, which displayed an atypical biochemical profile for the species (13).

Although there is abundant data showing the geographical distribution, host range, and infection sites of this pathogen, the mechanisms that lead to the establishment and progression of *H. ovis* infections remain unexplored. As shown in an invertebrate infection model, *H. ovis* strains originating from the uterus of dairy cows can display varying degrees of virulence (14). Based on the virulence phenotypes from that study, whole genome comparative analyses identified potential high virulence determinants in this organism, including Zinc ABC transporters, two hypothetical proteins, and a pathogenicity island (15). These comparative analyses created a blueprint for investigating *H. ovis* pathogenic capabilities, but the role these virulence factors may play in disease pathogenesis remains unexplored.

One of the most prevalent and costly animal diseases associated with *H. ovis* is metritis in dairy cows (5,16). This disease is characterized by acute uncontrolled opportunistic bacterial proliferation within the uterus in the face of immune dysregulation, leading to painful inflammation, impaired fertility, and sometimes death. The primary causative agents of metritis have not been clearly identified, but studies have shown that *Bacteroides pyogenes*, *Fusobacterium necrophorum*, *Porphyromonas levii*, and *Helcococcus ovis* are of importance in its etiology (5,17,18). Among these organisms, *H. ovis* is atypical in that it is a Gram-positive facultative anaerobic bacterium within an infection environment dominated by Gram-negative obligate anaerobes. Exploring the genomic features and diversity of this bacterium is a key step in unraveling its role in the pathogenesis of mixed infections.

Pangenomic analyses can provide valuable insights into the genomic diversity, virulence factors, and potential antimicrobial resistance profiles of bacterial populations. By examining the pangenome, which includes the core genome shared by all isolates and the accessory genome comprising genes unique to specific isolates, we can identify genetic variations that may be associated with virulence or adaptations to the uterine microenvironment in health or disease. A pangenomic analysis of *H. ovis* isolates from the uterus of healthy dairy cows and those with metritis can offer a comprehensive view of the bacterium’s genomic characteristics and shed light on its pathogenic potential, host adaptation, and implications for antimicrobial treatment options.

In this study, we sequenced the genomes of 35 *Helcococcus* clinical isolates from the uterus of dairy cows and tested their antimicrobial resistance and biochemical phenotypes. Phylogenetic and average nucleotide identity analyses placed 4 *Helcococcus* isolates within a cryptic clade, representing an undescribed species. We applied whole genome comparative analyses to explore the pangenome, resistome, virulome, and taxonomic diversity of the remaining 31 *H. ovis* isolates.

## Results

### *Helcococcus ovis* Isolation is Associated with Metritis

Of the thirty-eight cows evaluated for metritis diagnosis, twenty-one were healthy and seventeen were diagnosed with metritis. As shown in Table 1, six healthy cows and fifteen cows with metritis were culture-positive for *H. ovis*. We used Fisher’s exact test to examine the relationship between metritis and the presence of live *H. ovis* in the uterus, which showed a statistically significant association (p < 0.001). The odds ratio (OR) for cows with metritis was found to be 18.75 (95% confidence interval: 3.14-92.85), indicating an 18-fold increased risk of being culture-positive for *H. ovis* compared to healthy cows. Clinical data for the cows used for this study and for those associated with strains included from previous studies is presented in Supplemental File 1.

**Table 1.** Two-by-two contingency table of *H. ovis* isolation in healthy cows and cows with metritis.

|  | Healthy | Metritis | Row Sum |
| --- | --- | --- | --- |
| *H. ovis* Positive | 6 | 15 | 21 |
| *H. ovis* Negative | 15 | 2 | 17 |
| Column Sum | 21 | 17 |  |

$p < 0.001$ , OR=18.75 (95% CI: 3.14-92.85)

### Read Quality and Assemblies

A total of 30 *H. ovis* isolates were selected for Illumina sequencing. These included 20 recovered from the screening portion of this study and ten previously isolated strains. Illumina reads from two additional *H. ovis* strains (KG39 and KG40) were retrieved from the GenBank to make a total of 32 sets of Illumina reads. Two isolates (KG111 and KG115) were excluded from further analysis because they did not reach the desired coverage threshold of at least 15x. The remaining 30 samples had a mean of 163 mega base pairs (Mbp), ranging from 30 to 567. Using an expected *H. ovis* genome size of 1.8Mbp resulted in a mean coverage of 90x, ranging from 16x to 308x. Detailed read quality metrics and SRA accession numbers are listed in Supplemental File 2.

Of the resulting de-novo assemblies, five (KG101, KG116, KG93, KG118, KG97) resulted in unexpectedly small genome sizes ranging from 1.02 to 1.48 Mbp compared to the expected range of 1.7-.1.85 Mbp.

These genome assemblies also have fewer than the expected >1600 coding sequences (CDS) (1234–1537) and fewer than the expected 33 tRNAs (23–29) found in *H. ovis*, and therefore they were excluded from the pangenome analysis. However, they were retained for other analyses since they can provide valuable taxonomic and gene presence information. Finally, we also included five complete *H. ovis* genomes (KG36, KG37, KG38, KG104, and KG106) from a previous study (15). Genome assembly statistics for all thirty-five genomes included in this study and their accession numbers are listed in Supplemental File 3.

### *Helcococcus* Cryptic Strains

In a recent study we identified a single putative *H. ovis* strain (KG38) whose average nucleotide identity (ANI) with other *H. ovis* strains is lower than the suggested 96% threshold for same species determination (15). Although the initial identification of *H. ovis* isolates for this experiment was conducted based on 16S rRNA sequence identity comparisons, 16S rRNA sequence variations are often not specific enough to discriminate between closely related species (19). To screen for the presence of any cryptic species among our assembled genomes, we created a maximum likelihood phylogenetic tree and evaluated all-vs-all ANI relationships between all the strains in this study.

#### Taxonomy and ANI

As shown in Figure 1, four (KG38, KG95, KG105, and KG197) of the 35 strains included in this study cluster together in a clade forming an outgroup from the remaining 31 *H. ovis* strains. These strains have ANIs lower than 90% with the rest of the *H. ovis* species and also a higher than 96% ANI between each other. Although these three cryptic strains are closely related to *H. ovis,* their taxonomic position within the genus *Helcococcus* and family *Peptoniphilaceae* is unclear. We selected one representative *H. ovis* strain for each subclade within the species taxon and created a maximum likelihood phylogenetic tree which also includes the type strains for all species of the *Helcococcus* genus as well as the type species for the most closely related genera to *Helcococcus*: *Finegoldia* and *Parvimonas*. Figure 2 shows this phylogenetic tree alongside a heat map of ANI values between *H. ovis* strains and type strains of other close species and genera. Strains KG38, KG95, KG105, and KG197 form a cryptic clade within *Helcococcus*. Having less than 95% ANI with every other species of the *Helcococcus* genus is evidence that these strains represent a distinct novel species. Furthermore, having a higher than 70% ANI with the type strains of other *Helcococcus* species and less than 70% ANI with *Finegoldia magna* and *Parvimonas micra* is robust evidence that these strains belong to the genus *Helcococcus*. These observations are also supported by the maximum likelihood phylogenetic tree that was constructed with 232 orthologous genes shared across stains.

**Figure 1.**
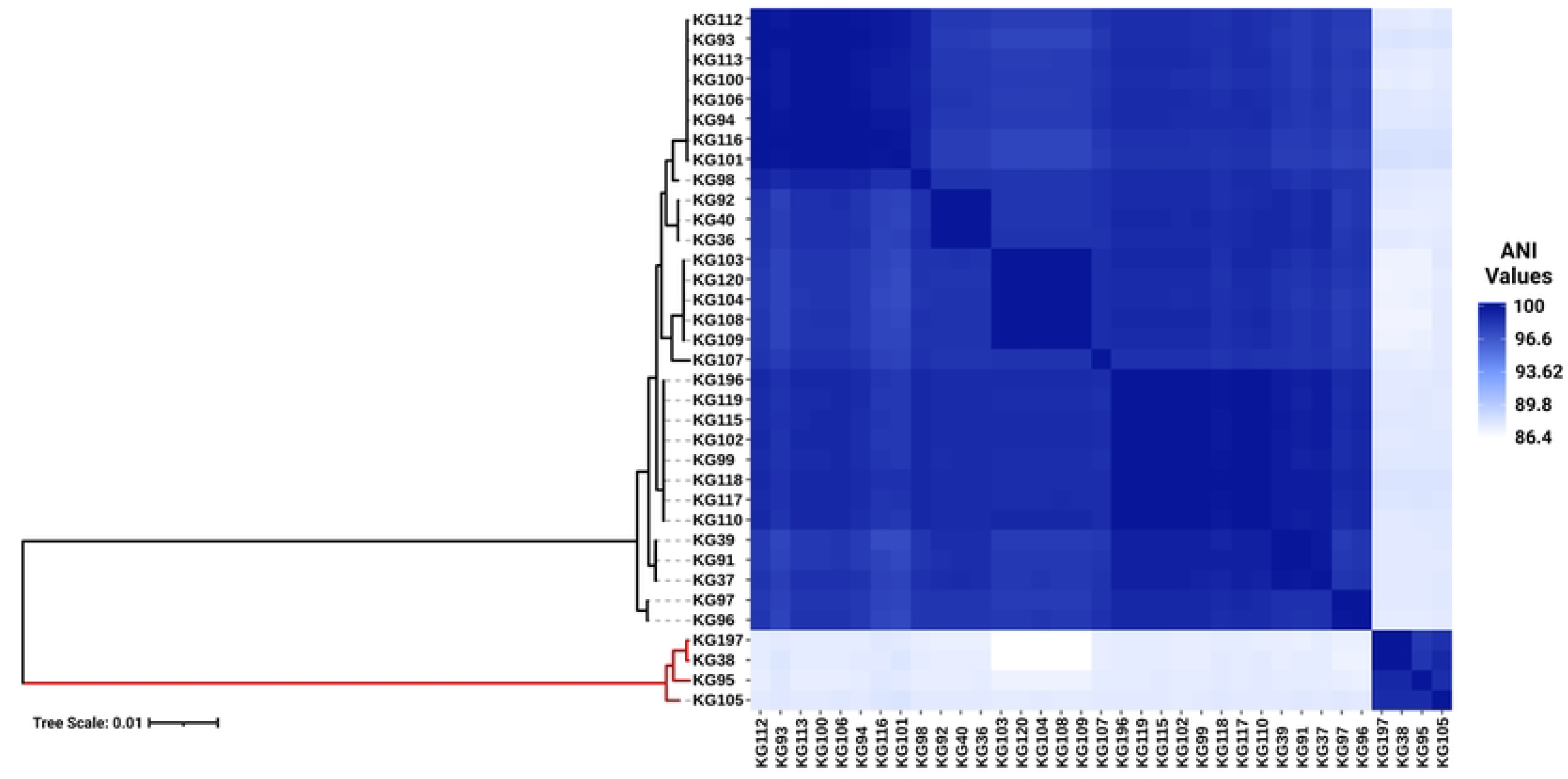
Heat map of whole genome average nucleotide identity based on BLAST+ (ANIb) and maximum-likelihood phylogenetic tree of 35 *Helcococcus* strains included in this study. The clade colored in red represents cryptic *Helcococcus* clade.

**Figure 2.**
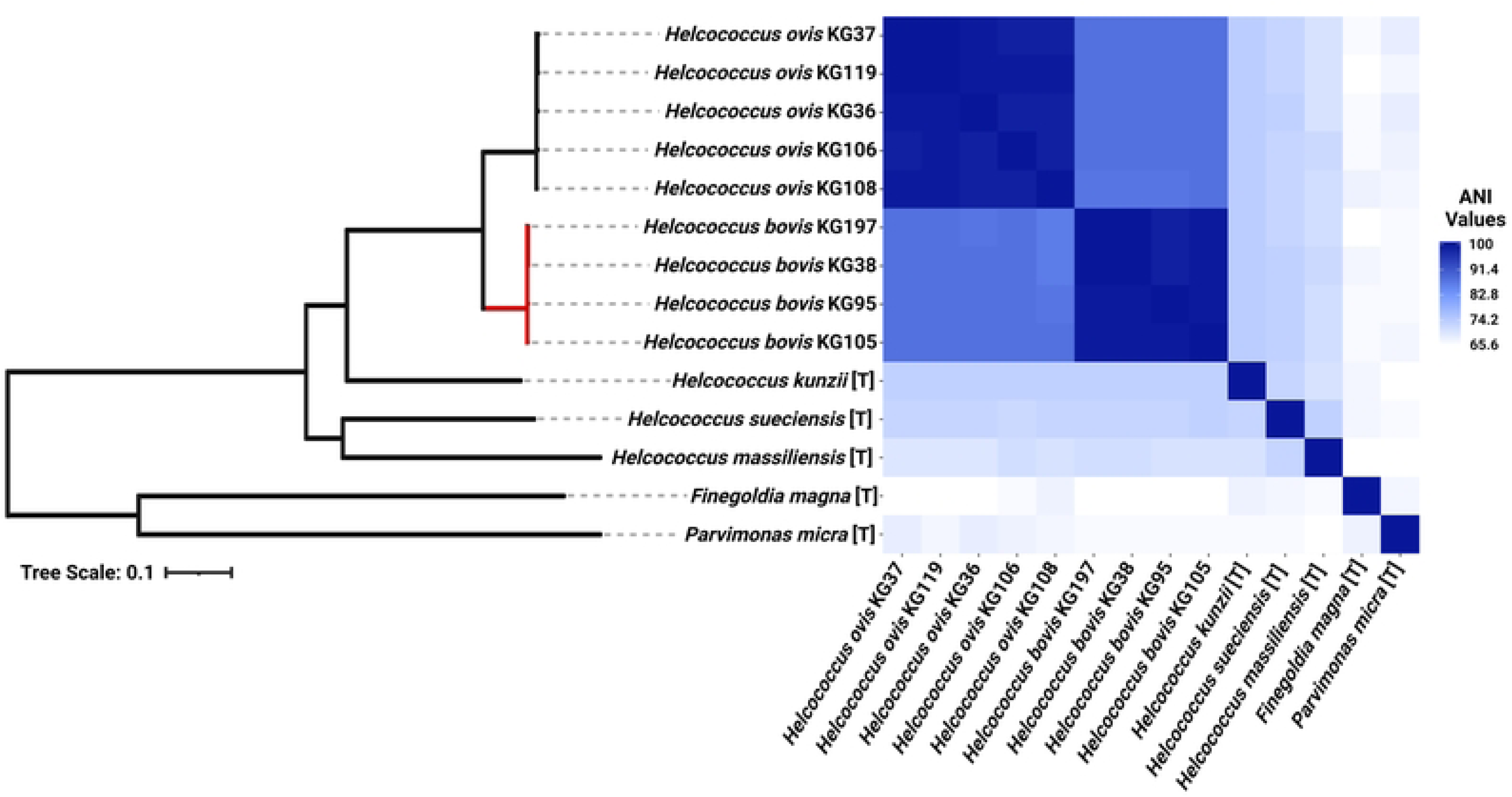
Heat map of whole genome average nucleotide identity based on BLAST+ (ANIb) and maximum-likelihood phylogenetic tree of the selected genomes of *Helcococcus ovis* and *Helcococcus bovis*, the type strains of the remaining species of the *Helcococcus* genus, and the type species of the most closely related genera to *Helcococcus*, *Finegoldia* and *Parvimonas*. [T] denotes the genome of a type organism.

Although the only publicly available whole genome sequences of *H. ovis* are from isolates associated with metritis in Holstein dairy cows, there are few publicly available near-complete *H. ovis* 16S rRNA sequences. As shown in Figure 3A, a multiple sequence alignment of near-complete *H. ovis* 16S rRNA sequences from this study, the Tongji strain, and the *H. ovis* type strain is able to discriminate between the core *H. ovis* clade and the cryptic strains. However, as shown in Figure 3A, these differences are driven by single nucleotide polymorphisms in hypervariable regions V2 and, to a lesser extent, V6. Based on this multiple sequence alignment, the *H. ovis* Tongji strain can also be considered a member of the cryptic *Helcococcus* sp. clade.

**Figure 3.**
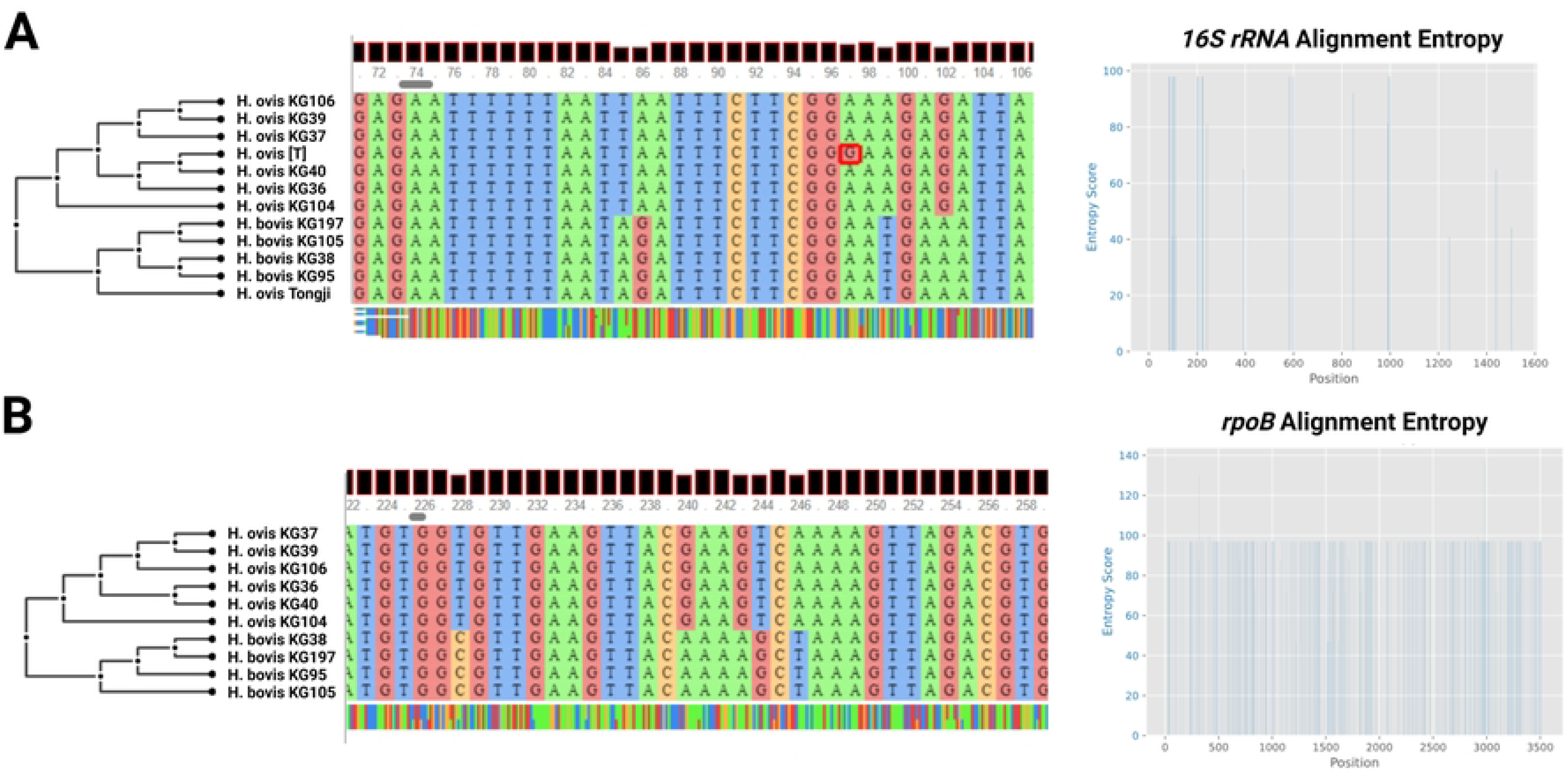
A) Multiple sequence alignment and sequence entropy plot of near-complete *Helcococcus ovis* 16S rRNA from this study, the *Helcococcus ovis* Tongji strain, and the *Helcococcus ovis* type strain. B) Multiple sequence alignment and sequence entropy plot of *rpoB* sequences from this study. Alignment windows display examples of areas of sequence entropy that can be used to differentiate between *Helcococcus ovis* and *Helcococcus bovis*.

In an attempt to identify a single marker gene to resolve the two *Helcococcus* sp. groups we also extracted the *rpoB* gene from the same genome assemblies in this study. Although there are no publicly available *rpoB* sequences for *H. ovis*, it is an often-used core gene for bacterial phylogenetic analyses. As shown in Figure 3B, the multiple sequence alignment is able to discriminate between the core *H. ovis* clade and the cryptic strains while also having areas of sequence entropy across the gene, making it a better candidate single marker gene than 16S rRNA for Helcococcus species differentiation.

#### Proteome comparison

Finally, as shown in Figure 4, a protein blast alignment between the complete proteomes of three representative *H. ovis* strains (KG36, KG37, and KG106) and three of the cryptic strains (KG38, KG95, and KG105) illustrates that the cryptic strains have protein sequence identities with the reference *H. ovis* KG36 as low as 80-70% across their genomes.

**Figure 4.**
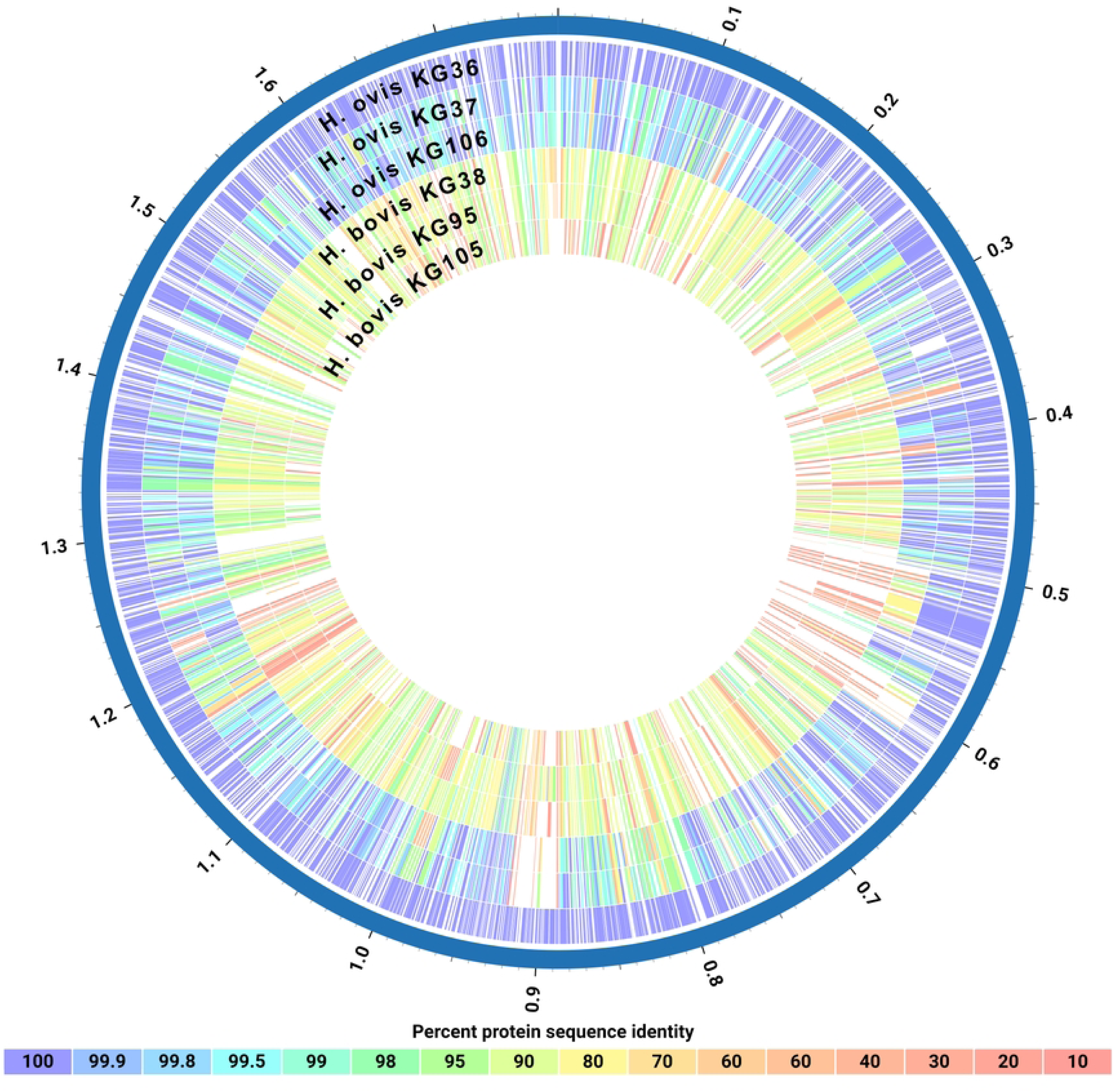
Circos plot of protein sequence alignments of three *Helcococcus ovis* and three *Helcococcus bovis* strains. Percent protein sequence identities were calculated against the proteome of reference strain *Helcococcus ovis* KG36.

#### Phenotypic characteristics

Phenotypically, the cryptic *Helcococcus sp.* strains are Gram-positive facultative anaerobic cocci that depend on pyridoxine supplementation for growth in vitro. They can be cultivated at 36-38 °C on tryptone soy agar (TSA) with 5% defibrinated sheep blood and 0.001% pyridoxal HCl. After 72-96 hours of incubation, they form pinpoint transparent colonies morphologically indistinguishable from *H. ovis* and displaying little to no alpha hemolysis. As shown in Figure 5, cryptic strain KG38 displays weak hemolysis on blood agar when compared to *H. ovis* strains. In liquid medium, both *H. ovis* and the cryptic strains grow well in brain heart infusion (BHI) broth supplemented with 0.1% Tween80 and 0.001% pyridoxal HCl.

**Figure 5.**
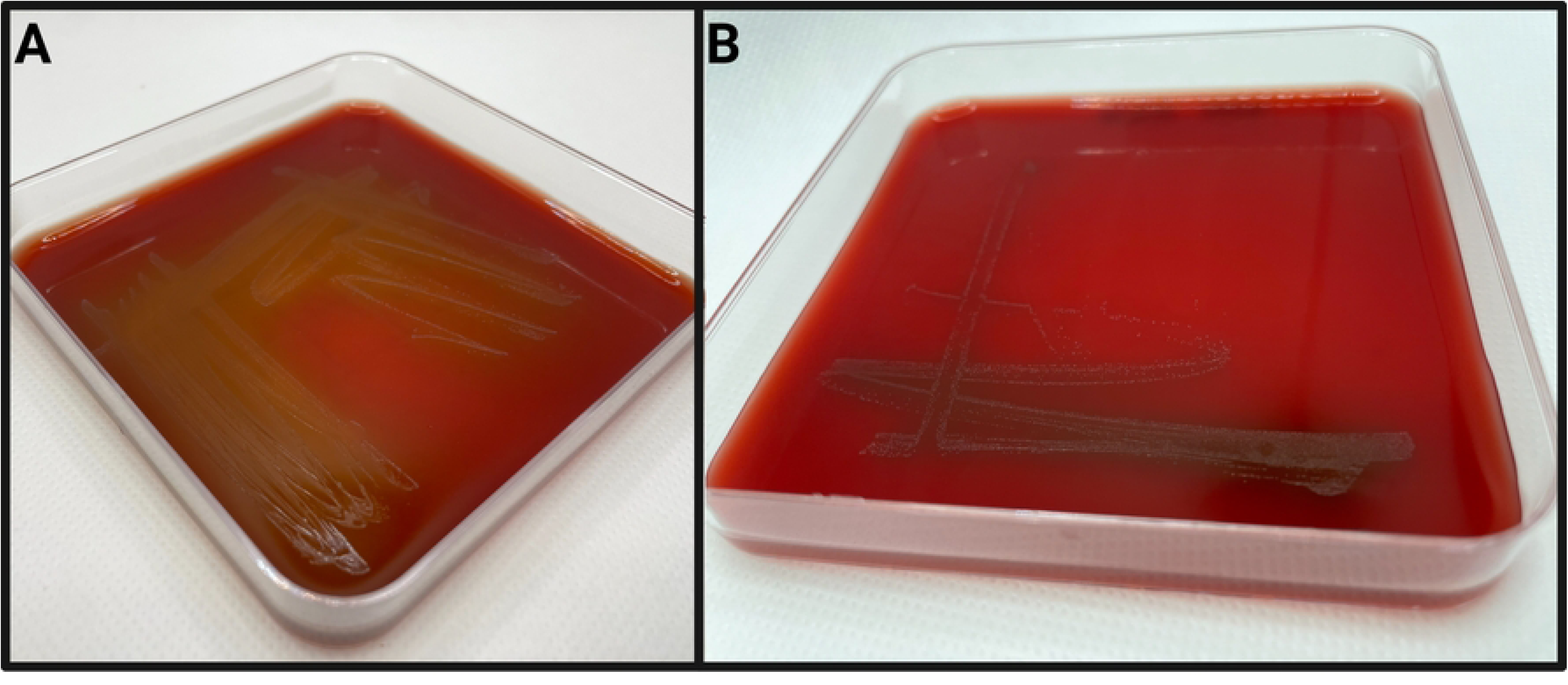
Examples of *Helcococcus ovis* KG37 (A) and *Helcococcus bovis* KG38 (B) culture on tryptone soy agar with 5% defibrinated sheep’s blood and 0.001% pyridoxal HCl after 72 hours of incubation.

As shown in Table 2, eight isolates were assessed to identify their enzymatic activity. *H. ovis* strains KG36, KG37, KG104, and KG106, and cryptic strains KG38, KG95, 105, and KG197 exhibited positive results for both alanine arylamidase and L-proline arylamidase. However, in contrast to the cryptic strains, *H. ovis* strains also demonstrated positive results for at least one of the following: tyrosine arylamidase (3/4), Beta galactopyranosidase (2/4), D-mannose (3/4), or D-maltose (1/4). Cryptic strain KG38 was the sole strain positive for leucine arylamidase and alanyl-phenylalanyl-proline arylamidase. Although the Vitek 2 Gram-Positive ID card is capable of identifying *Helcococcus kunzii* based on its biochemical profile, it is not designed to identify *H. ovis*. As a result, all but one tested strain produced "low confidence" or "unknown" species identification. Cryptic strain KG38 was misidentified as 99% probability "*Dermacoccus nishinomiyaensis*/*Kytococcus sedentarius*". These results indicate that differentiating between *H. ovis* and the cryptic *Helcococcus* strains may be possible based on the absence of specific enzymatic activity beyond alanine arylamidase and L-proline arylamidase. However, a larger sample size is needed to draw any conclusions regarding the differential enzymatic activities of these two clades.

**Table 2.** Biochemical characteristics and Vitek2 identification results of *Helcococcus sp.* isolates. Only tests with at least one positive result are included. A complete list of biochemical tests is presented in Supplemental File 4.

| Isolate | APPA | LeuA | AlaA | ProA | TyrA | BGAR | dMAN | dMAL | Vitek2 ID |
| --- | --- | --- | --- | --- | --- | --- | --- | --- | --- |
| *Helcococcus ovis* KG104 | - | - | + | + | + | + | - | - | Low Discrimination |
| *Helcococcus ovis* KG106 | - | - | + | + | + | - | + | - | Low Discrimination |
| *Helcococcus ovis* KG36 | - | - | + | + | - | + | + | - | Unidentified |
| *Helcococcus ovis* KG37 | - | - | + | + | + | - | + | + | *Granulicatella elegans*<br>(95% Probability) |
| *Helcococcus bovis* KG38 | + | + | + | + | - | - | - | - | *Dermaococcus nishinomiyaensis/Kytococcus sedentarius*<br>(99% Probability) |
| *Helcococcus bovis* KG95 | - | - | + | + | - | - | - | - | Low Discrimination |
| *Helcococcus*<br>*bovis* KG105 | - | - | + | + | - | - | - | - | Low Discrimination |
| *Helcococcus*<br>*bovis* KG197 | - | - | + | + | - | - | - | - | Low Discrimination |

### *H. ovis* Pangenome Construction and Associations

The four genomes belonging to the cryptic strains were excluded from pangenomic analyses as this study aims to explore the pangenome of *H. ovis*, the *Helcococcus* species associated with metritis in dairy cows. A total of 31 genomes were initially included in the pangenome construction. However, there were low-quality assemblies that did not result in complete enough genomes to warrant inclusion into the pangenome. As shown in Figure 6, the number of new genes in the pan-genome plateaus after 25 genomes are included. We therefore excluded the five lowest-quality genome assemblies from the pangenome construction and retained 26 assemblies in the analysis. The exclusion of these low-quality assemblies resulted in the core genome expanding from 683 to 1045 gene families. The resulting core, soft core, shell, and cloud genomes are shown in Figure 7. In short, the *H. ovis* pangenome consists of 845 core genes (99% <= strains <= 100%), 203 soft core genes (95% <= strains < 99%), 1078 shell genes (15% <= strains < 95%), and 556 cloud genes (0% <= strains < 15%).

**Figure 6.**
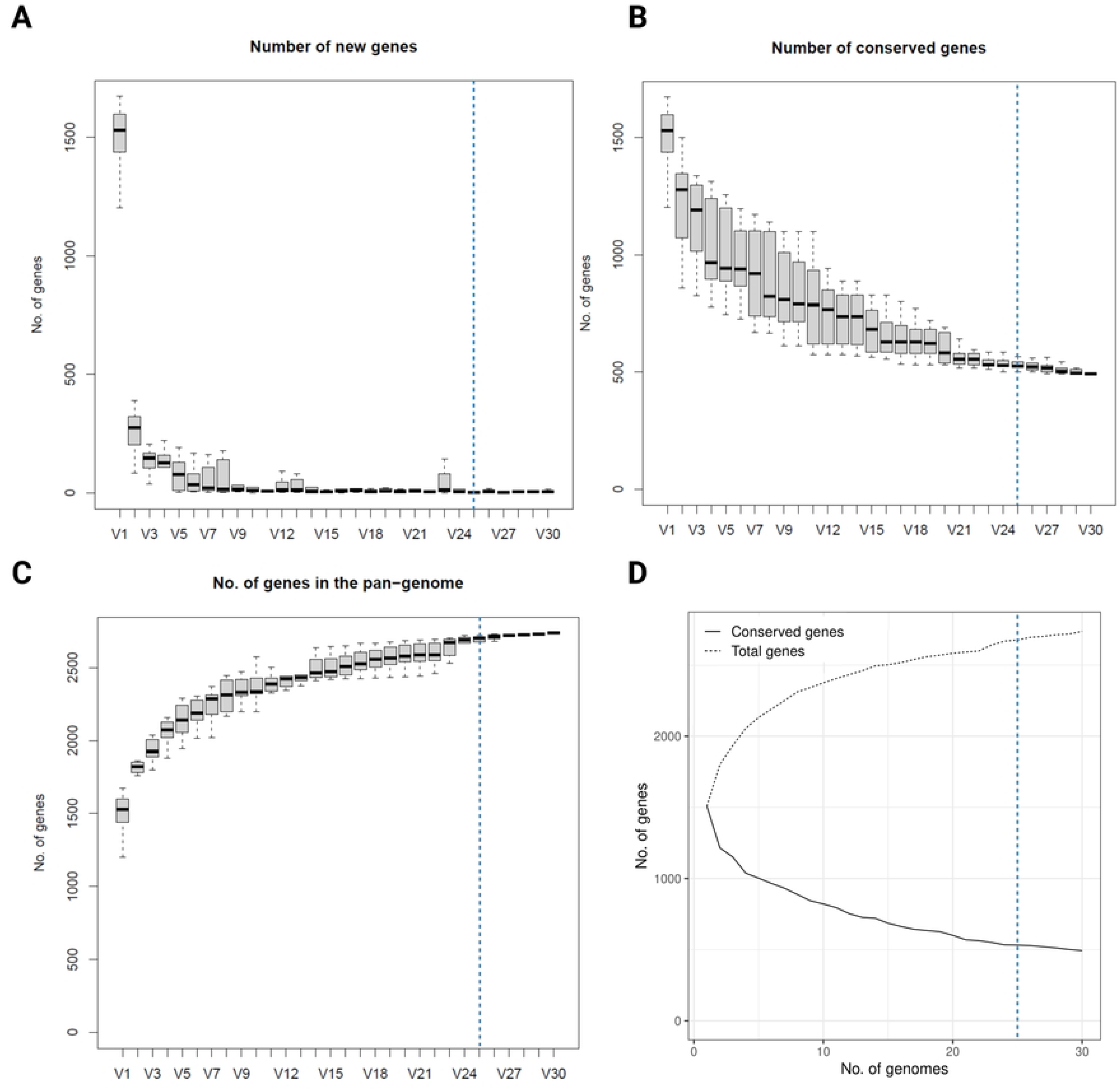
Plots depicting how the pangenome varies as genomes are randomly added to the pangenome construction. The dashed blue line marks the 25-genome threshold selected for this study.

**Figure 7.**
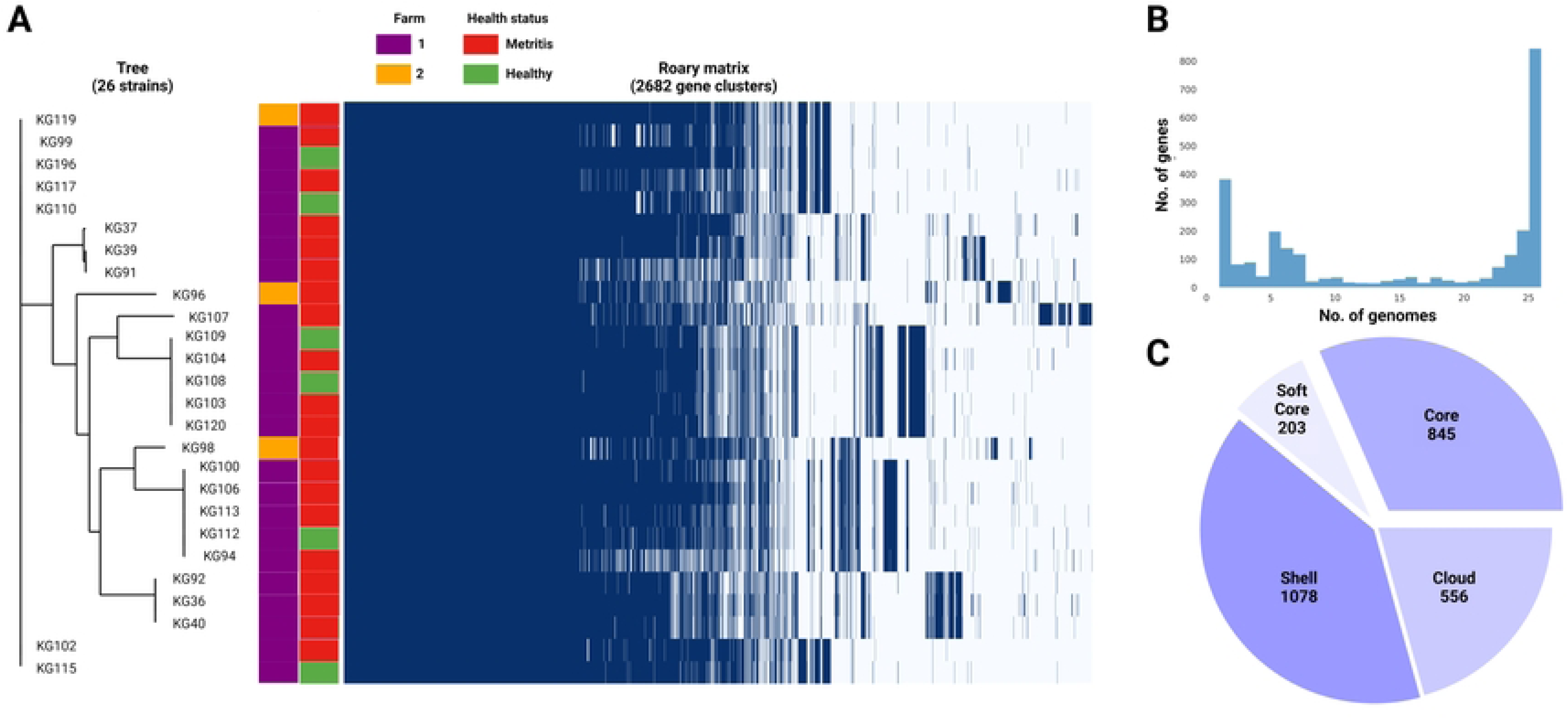
A) *Helcococcus ovis* pangenome gene matrix depicting the 2682 gene clusters identified by Roary. B) Pangenome frequency plot depicting how many gene clusters are found in only 1 to only 25 genomes. C) Pie chart summarizing the pangenome structure.

The final *H. ovis* pangenome includes 20 strains from cows with metritis and six strains from healthy cows. The complete *H. ovis* pangenome, including the gene presence and absence table, is presented in Supplemental File 5. We were interested in finding genes that were enriched in the metritis or healthy group of strains. We ran a Scoary (20) analysis using both the 31-strain and the 26-strain pangenomes with the Benjamini-Hochberg adjusted p-values to identify the genes most overrepresented in a specific host group. Using a significance level of p<0.05 we did not find any gene group overrepresented in either of the host groups.

### Virulome

After using Abricate for mass screening of virulence factors in the 26 *H. ovis* genomes against the Virulence Factor Database, no positive hits were returned. To further investigate the virulome of 26 *H. ovis* strains in this study, we curated a set of 22 putative virulence factor genes based on previous comparative genome analyses (15). The resulting virulome is presented in Figure 8 as a heat map. There is no observable pattern in the presence or absence of virulence factors in these strains in relation to the health status of the host or farm location.

**Figure 8.**
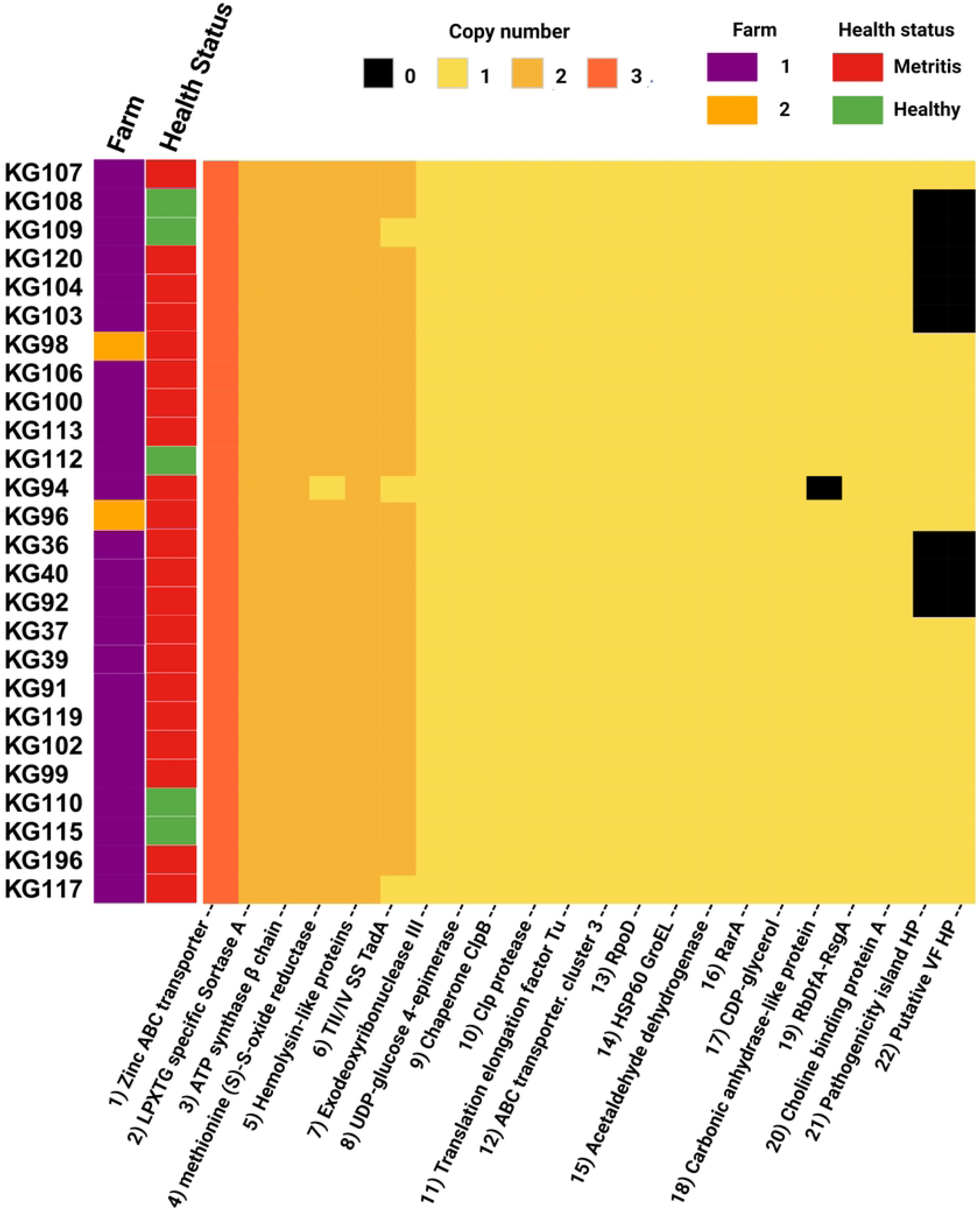
Heat map of virulence factors of *Helcococcus ovis* strains explored in this study.

Two hypothetical proteins and a pathogenicity island have been recognized as potential high virulence determinants of *H. ovis* in invertebrate infection models (14). These high virulence determinant CDS are found in 69% (18) of the strains in this study and are absent in only eight strains. Both the hypothetical proteins and the pathogenicity island are co-occurring in every genome where they are present and are altogether absent in the remaining strains. We used mauve to visually inspect the spatial distribution of these co-occurring high virulence determinants in the two complete *H. ovis* genomes where they are present. In both KG37 and KG106, the two hypothetical protein CDS are found closely associated with a ZnuABC locus located more than 500,000 base pairs away from the co-occurring pathogenicity island. To further explore the cause of the co-occurrence and co-absence of these virulence determinant CDS we identified and excluded loci containing elevated densities of base substitutions in the twenty-six genomes and built an approximately-maximum-likelihood phylogenetic tree (Supplemental File 6). The clades that do not contain the high virulence determinant CDS seem to be spread across the tree, showing that the co-occurring CDS are not restricted to a single lineage but are found in multiple, more distantly related lineages.

### Resistome and Plasmids

We also used Abricate for mass screening of acquired antimicrobial resistance genes (ARGs) against the Comprehensive Antibiotic Resistance Database. The search was limited to acquired resistance genes because not enough experimental data is available for *H. ovis* to evaluate resistance-associated point mutations. We also screened for known plasmid sequences by querying against the PlasmidFinder database. The results of these analyses are presented in Figure 9.

**Figure 9.**
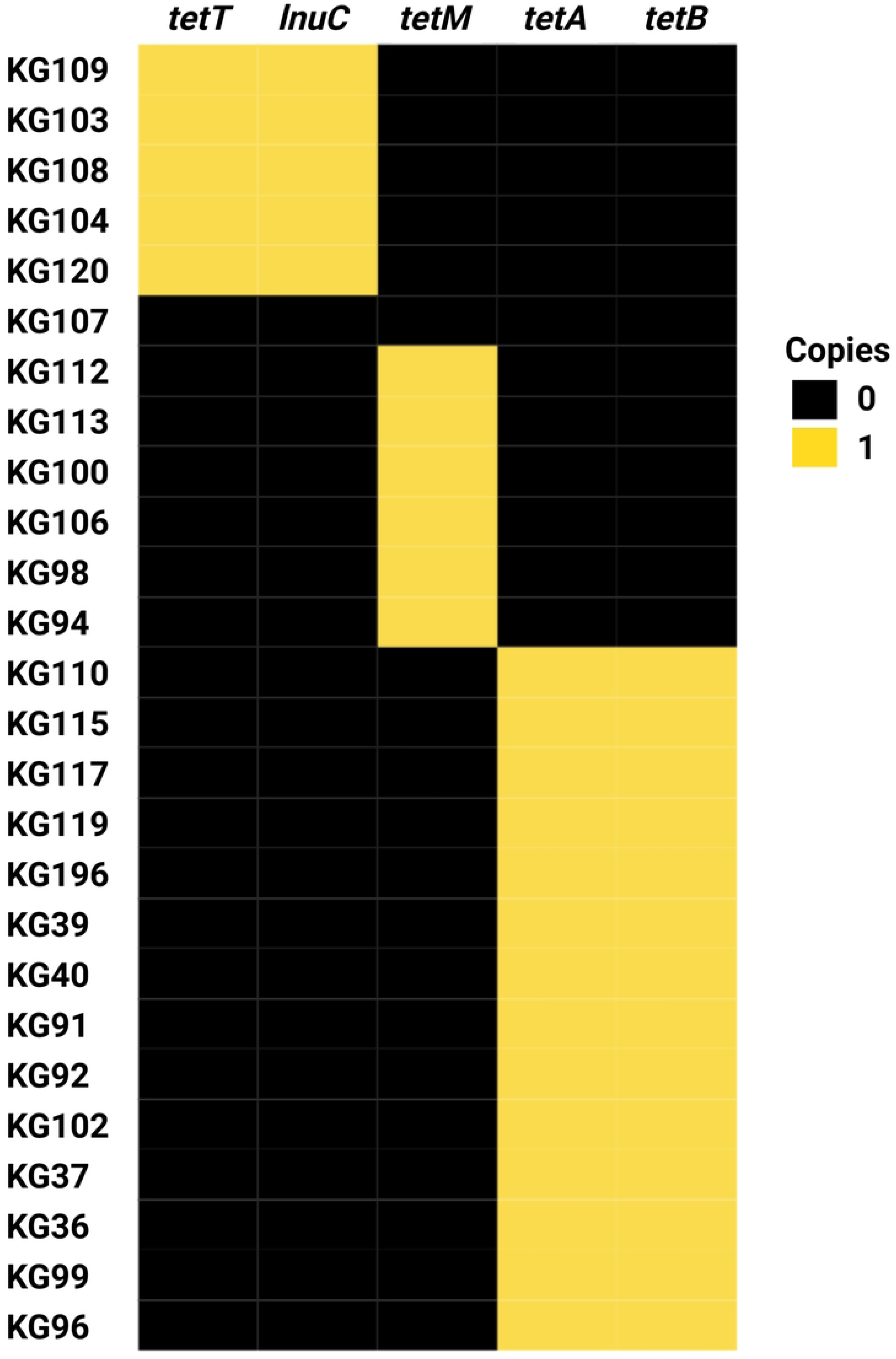
Heat map of antimicrobial resistance gene profiles of *Helcococcus ovis* strains sequenced in this study.

Strain KG107 is the only one of the thirty screened genomes that does not contain any acquired ARGs. Nine *H. ovis* strains carry only *tetM*, 15 strains carry both *tetA* and *tetB*, and five strains carry both *tetT* and *lnuc*. With the exception of *lnuc,* which confers resistance to lincosamides, all other ARGs found in this experiment confer resistance to tetracyclines. Acquired antimicrobial resistance genes *tetA* and *tetB* are, in all strains, located within a prophage region commonly found within *H. ovis* genomes. Similarly, *tetT* and *lnuC* are found in conjunction within a prophage region in all strains where they occur. This suggests prophage integration events are a significant driver of ARGs acquisition in *H. ovis*. Alternatively, *tetM* is located within a previously described integrated plasmid region (repUS43_1_CDS12738(DOp1)), often found in *Streptococcus spp*.

A total of ten strains were selected to be evaluated for resistance to 22 clinically relevant antimicrobials. Subsets of strains from each AMR genotype including *tetM* only (KG100, KG106, and KG113), *tetA*/*tetB*(KG36, KG37, KG92, KG196), *tetT*/*lnuC* (KG104, KG109, KG120), and none (KG107), were selected for minimum inhibitory concentration (MIC) testing. As shown in Table 3, MICs are reported in μg/mL without antimicrobial resistance breakpoint interpretations because there are currently no interpretive standards established by the Clinical and Laboratory Standards Institute (CLSI) for *H. ovis* in the uterus of cattle. We used the MIC results of strain KG107 as the wild-type reference since it was the only isolate that does not carry any known ARG. Strains carrying any tetracycline resistance gene (*tetT, tetM*, or *tetA*/*tetB*) had a higher MIC for tetracycline. The wild-type strain had a tetracycline MIC ≤ 0.250 μg/mL, and all other strains had a tetracycline MIC ≥ 1.0 μg/mL. Although strains carrying *tetM* or *tetT* displayed resistance to doxycycline and minocycline compared to the wild type, strains carrying *tetA*/*tetB* did not follow the same pattern. None of the *tetA*/*tetB* positive strains had increased resistance to minocycline, and their resistance to doxycycline was inconsistent and less than that of *tetM* and *tetT* positive strains. Although MIC for lincomycin were not evaluated, the strains that carry *lnuC* (KG104, KG109, KG120) did not show resistance to clindamycin, the only lincosamide tested.

**Table 3.**
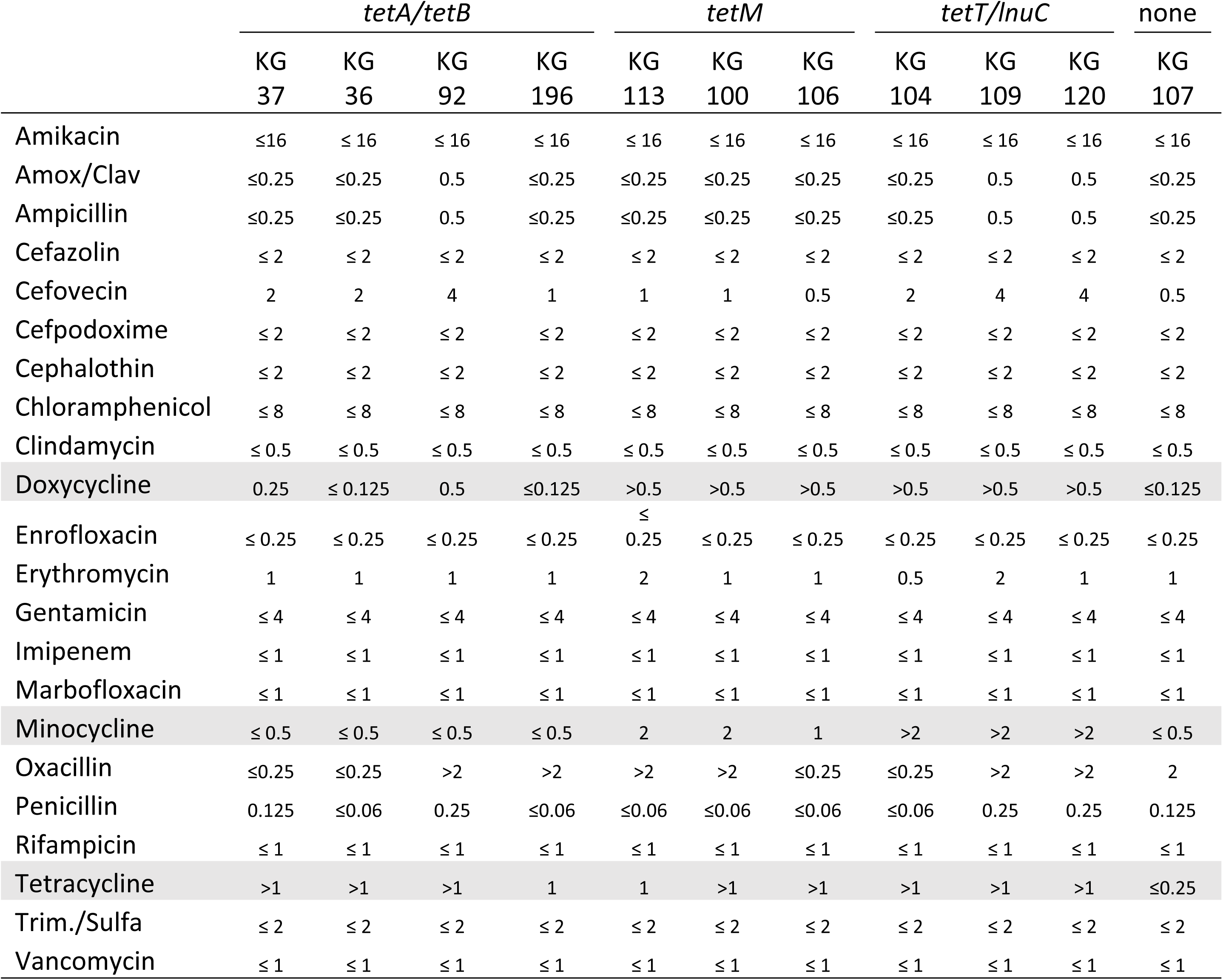
Antimicrobial resistance profiles of *Helcococcus sp.* Minimum inhibitory concentration values are in microgram/milliliter (μg/ml). The rows of tetracycline class antibiotics are highlighted in grey.

Finally, as shown in Table 3, some strains showed resistance to cefovecin and oxacillin without carrying any ARG known for conferring resistance to beta-lactams. We ran a Scoary (Brynildsrud et al., 2016) analysis using a resistance threshold of ≥ 2 for cefovecin and ≥ 0.5 for oxacillin, using the CLSI soft-tissue cutoffs for non-*Staphylococcus aureus* staphylococci in dogs and cats as an approximation (21). We used the Benjamini-Hochberg adjusted p-values to identify the genes most overrepresented in a specific host group. We did not find any gene group overrepresented in either of the host groups using a significance level of p<0.05. This suggests that beta-lactam resistance in *H. ovis* may be mediated by chromosomal mutation resistance instead of mobile ARGs.

## Discussion

In this study, we examined the genomes of *H. ovis* strains obtained from the uteri of both healthy dairy cows and those with metritis. Our analysis focused on exploring the pangenome, resistome, virulome, and taxonomic diversity of these strains. Additionally, we sought bacterial genome-wide associations between *H. ovis* gene clusters and metritis in dairy cows.

While the costs of second-generation short-read whole-genome sequencing (WGS) have significantly decreased in the past decade, third-generation long-read sequencing remains a less affordable emerging technology. In our study, we opted for the more commonly used approach of employing low-depth Illumina short-read sequencing to maximize the inclusion of a larger number of strains in our analysis. As is evident in our findings, this approach may lead to the loss of genomes due to low coverage and incomplete assemblies. However, this tradeoff is acceptable when considering the low-cost, high-throughput generation of excellent-quality reads (Q>35).

The *Helcococcus* genus is comprised of five well-described species. However, their genetic diversity remains unexplored due to the limited availability of sequenced genomes. Within this small genus there are still unresolved and contradictory taxonomic classifications. For example, the species *H. pyogenes* was described as a new species isolated from a prosthetic joint infection based on biochemical tests and a partial (518bp) 16S rRNA sequence identity in 2004 (2). A later study proposed another new species, *H. seattlensis*, isolated from a human with urosepsis also based on 1512bp 16S rRNA sequence identity, which also shares 99.4% sequence identity with *H. pyogenes* suggesting it is likely the same species (3). Another taxonomic uncertainty within the genus is the classification of the *H. ovis* Tongji strain, isolated from the only recorded *H. ovis* infection in a human (13). This strain displayed an atypical biochemical profile for *H. ovis* and a 98.9% 16S rRNA sequence identity with the *H. ovis* type strain, which led researchers to question its place within the species taxon. Phylogenomic analyses have shown that, based on 16S rRNA sequence identity, the Tongji strain belongs to a subclade of the species also populated by *H. ovis* strain KG38 (15). Whole genome-based multi-locus phylogenomic analyses in this study confirmed these findings and identified three further strains (KG95, KG105, and KG197) belonging to the cryptic clade. Average nucleotide identity, proteome identity, and phenotypic analyses provide robust evidence that these strains belong to a distinct novel species, for which we propose the name *H. bovis* sp. nov. (bo’vis. L. gen. n. bovis of the cow). *H. ovis* and *H. bovis* strains share 87-89% average nucleotide identity with each other placing their relationship in the 0.2% of pairs that fall within the 83– 95% ANI valley range (22). This makes the relationship between the two species a rare candidate for exploring bacterial speciation mechanisms and the role horizontal gene transfer has on the speciation process. These *H. bovis* strains originated from two geographically separate farms in North Central Florida. They were also retrieved from uteri of both cows with metritis (1) and healthy cows (3).

However, the sample size in this study is too small to draw conclusions about the association between the presence of *H. bovis* and uterine health status. Strain KG38, part of the novel species group, has been shown to have attenuated virulence when compared to other *H. ovis* isolates (14). Since *H. bovis* occupies a similar biochemical niche as the more virulent *H. ovis* strains, its role as a commensal organism of the reproductive tract is an area that warrants further exploration. Although the multiple sequence alignment of the whole 16 S rRNA gene can discriminate between *H. ovis* and *H. bovis*, the responsible sequence variations are in hypervariable regions V2 and V6 which are not often targeted in metagenomic amplicon studies. This means sequences belonging to *H. bovis* will contribute reads to the amplicon sequence variants classified as *H. ovis* in most metagenomic studies amplifying the V3-V4 hypervariable regions. Unlike the 16S rRNA sequence, *rpoB* has more regions of dissimilarity between *H. ovis* and *H. bovis*, making it a much more useful single-marker gene to resolve these two *Helcococcus* species.

We screened a subset of cows from this study to show that isolation of *H. ovis* from the uterus of dairy cows is strongly associated with metritis. Previous studies have shown that *H. ovis* DNA is more abundant in relative and absolute terms in the uterus of dairy cows with metritis than in healthy cows at the time of metritis diagnosis (5,17). Although all healthy cows have been shown to harbor *H. ovis* genomic DNA (gDNA) in the uterus after parturition, this gDNA is not indicative of the presence of viable bacteria, likely because healthy cows are able to mount adequate immune responses that neutralize these organisms (23,24). Previous to this study, isolation of live *H. ovis* from the uterus of dairy cows had been limited to only cows with metritis, and targeted comparative cultivation screenings have not been conducted (10,25). These results are a robust addition to the current body of evidence showing *H. ovis* is one of the key organisms in the pathogenesis of metritis in dairy cows at the time of diagnosis.

Our inability to find any *H. ovis* genotype association with metritis is likely due to the fact that metritis is characterized by a dysbiosis of the uterine microbiota that is unlikely to be explained by gene groups within a single component bacterial species (18,26). Furthermore, as is the case in microbial communities in gut dysbiosis, it is possible that *H. ovis* is more prevalent in the diseased uterus because the disease condition widens an independent metabolic niche which the bacterium can then fill without having to play a key role in the necessary steps for the development of disease (27). The genome-wide association analyses conducted in this study have been successfully used to find the genetic basis for high penetrance phenotypes in bacteria like virulence (28) or antimicrobial resistance (29), but we were not able to establish any phenotype-genotype link with this approach. In this study we also measured simple phenotypes like hemolysis and pyridoxal dependence in vitro for all strains but did not find any insightful phenotypic variation between them.

Since *H. ovis* is not a well-known pathogen or a model organism, there is no experimentally verified virulence factor (VF) data. We found that the putative pathogenicity island and the hypothetical VF associated with a Zinc ABC transporter locus were either co-occurring or altogether absent from *H. ovis* strains. Manual screening of the available complete genomes revealed that these putative VFs are not part of the same operon in the species. This raises the question of whether they are functionally linked or if they display this pattern in our samples by chance. As shown in Supplemental File 6, these putative high virulence determinants are present and absent across different subclades of *H. ovis* and do not show evidence of being driven by the founder effect. According to ProteInfer (30), both the ZincABC transporter-associated VF and the conserved membrane-spanning protein within the pathogenicity island participate in metal ion binding and transport. This suggests that, as has been shown in *Escherichia coli* (31), these *H. ovis* accessory virulence genes co-occur due to having connected functions, and the resulting phylogenetic distributions are the result of convergent gene loss instead of founder effect.

All the ARGs found in our strains are located within mobile genetic elements like plasmids or prophage regions, which makes *H. ovis* a reservoir of mobile ARGs in a food production setting. All but one *H. ovis* strain (KG107) contain ARGs conferring resistance to tetracyclines. This strain is a valuable clinical isolate since it can be used as a wild-type reference strain with no ARGs to benchmark the susceptibility of *H. ovis* to antimicrobials. Unlike *tetA*/*tetB* positive strains, strains carrying genes encoding the cytoplasmic ribosomal protection proteins TetM or TetT, also have elevated resistance to doxycycline and minocycline. Oxytetracycline and ceftiofur are the only two antimicrobials labeled in the United States for the treatment of metritis in lactating dairy cattle. However, the United States Food and Drug Administration has banned the extra-label use of ceftiofur in animals, and may move towards policies like the Netherlands where the use of ceftiofur administration to agricultural animals is restricted (32). The alternative, intrauterine oxytetracycline infusions, remains a frequent practice both in clinical and research settings in the United States and Europe (33,34). Furthermore, dairy operations often use oxytetracycline as prophylactics in heifer rearing or as a treatment for calf pneumonia, a type of infection *H. ovis* has been implicated in (12). Although the MICs for oxytetracycline were not assessed, there is no inherent difference between a tetracycline and an oxytetracycline resistance genes (35), and 96.8% of isolates in this study carry at least one tetracycline resistance gene shown to confer tetracycline resistance in vitro. These findings indicate that further studies are needed to evaluate the effectiveness of tetracyclines as a treatment for metritis in dairy cattle.

## Conclusion

This study found that the presence of viable *H. ovis* in the uterus of dairy cows is associated with metritis. However, we found no evidence that a specific *H. ovis* genotype or gene cluster is associated with the disease. Virulence factor comparisons showed two putative high virulence determinants are common but have varying prevalence in these strains with a phylogenetic distribution consistent with convergent gene loss. Based on the genetic dissimilarity and phenotypic characteristics, strains KG38, KG95, KG105 and KG197 represent a novel species of the genus *Helcococcus*, for which we propose the name *Helcococcus bovis* sp. Nov. (bo’vis. L. gen. n. bovis of the cow). The type strain for this species is KG38 (Accession number CP121192). The significance of this species in the context of uterine health remains to be explored. The majority (30/31) of *H. ovis* strains in this study carry antimicrobial resistance genes conferring resistance to tetracyclines, which has significant clinical consequences for the treatment of metritis and other *H. ovis-*associated respiratory infections in cattle. The convergence of widespread ARG-mediated tetracycline resistance in these uterine pathogens and initiatives to phase out the use of ceftiofur to treat metritis reveals an immediate need to find alternative treatments and prevention strategies for this important animal disease.

## Materials and Methods

### Metritis Diagnosis and Uterine Sample Collection

All procedures involving cows were approved by the Institutional Animal Care and Use Committee of the University of Florida; protocol number 201910623. In this study, a total of 43 lactating Holstein Friesian cows were used. Three cows were from North Florida Holsteins and 40 were from the University of Florida’s Dairy Research Unit, both located in north central Florida. Each cow had uterine discharge collected directly from the uterus with a sterile pipette, and evaluated at four, six, and eight days postpartum. 500uL of uterine discharge was suspended in 500uL of BHI broth with 30% glycerol and stored at -80 °C.

The uterine discharge was scored on a 5-point scale (Jeon et al., 2016). Score 1 indicates normal lochia, viscous, clear, red, or brown discharge that was not fetid; Score 2 indicates cloudy mucoid discharge with flecks of pus; Score 3 indicates mucopurulent discharge that was not fetid with less than 50% pus; Score 4 indicates mucopurulent discharge that was not fetid with more than 50% pus; and Score 5 indicates fetid red-brownish, watery discharge. Cows with uterine discharge scores of 1-4 were considered healthy, whereas those with a score of 5 were diagnosed with metritis. Nine cows that had a uterine discharge score of 1 to 4 at the time of sampling but developed metritis sometime in the 21 days after parturition were excluded from microbiological testing.

### Bacteria Isolation and Identification

To selectively culture *H. ovis* from uterine discharge samples, 20μL of the discharge suspension was streaked onto *Helcococcus* selective agar. The agar plates were incubated for 72 hours at 36°C under aerobic conditions with 6% CO2 (11). Following incubation, individual pinpoint nonpigmented colonies were selected and sub-cultured on tryptone soy agar with 5% defibrinated sheep blood and 0.001% pyridoxal HCl for propagation. Species determination of the isolates was performed via comparative analysis of Sanger sequencing of their 16S rRNA genes.

### Whole Genome Sequencing

Genomic DNA (gDNA) was extracted using the DNeasy blood and tissue kit following the manufacturer’s instructions (Qiagen). Genomic DNA purity was measured using a NanoDrop 2000 spectrophotometer; final DNA concentration was confirmed with a Qubit 2.0 Fluorometer. DNA integrity was visualized via agarose gel electrophoresis. Library preparation was done with the Nextera XT kit (Illumina, Inc.), following the manufacturer’s instructions, and it was loaded into the MiSeq reagent kit V2. Sequencing was performed on a MiSeq platform (Illumina, Inc.) with a 2 × 250-bp 500-cycle cartridge. Seven previously sequenced strains were also included in this study. Two of them consist of Illumina sequenced draft genomes KG39 (Accession number SRX5460741) and KG40 (Accession number SRX5460742). The remaining five are complete genomes that were hybrid assembled using ONT and Illumina sequencing for genomic comparisons performed in a previous study (15) (KG36, KG36, KG38, KG104, KG106).

### Genome Assembly and Annotation

After performing quality control with fastp (36) the resulting reads were evaluated using MultiQC (v1.14).The minimum coverage threshold for inclusion in the study was set at 15x (37). De-novo genome assembly was performed using Unicycler (v0.5.0) (38). Assembly quality was assessed using Benchmarking Universal Single-Copy Orthologs (v4.1.2) (39). Genome annotations were conducted using Prokka and the genome annotation service in BV-BRC using the RAST tool kit (40,41)

### Taxonomic analyses

Whole genomes of *Helcococcus spp.* and the type strains of the recognized species within the genera *Helcococcus, Finegoldia, and Parvimonas* were used to create a phylogenetic tree with the BV-BRC codon tree pipeline using 500 single-copy PGFams (42). In order to verify that the constructed phylogenetic tree was not affected by recombination events, we used Snippy (v4.6.0) to align Illumina reads of the 26 *H. ovis* genomes using the *H. ovis* KG37 complete genome assembly as reference (43). We then used Gubbins (v3.3.3) to identify loci affected by recombination and construct a phylogeny based on point mutations outside of these regions (44). Phylogenetic trees were visualized and annotated using Interactive Tree of Life (iTOL v5) webtool (45). Average nucleotide identities (ANI) were calculated via BLAST pair-wise comparisons of all sequences shared between two strains (ANIb) using the JSpecies web server (Richter et al., 2016). 16S rRNA gene sequences were extracted from the raw paired-end Illumina reads using phyloFlash (v3.4.2) and *rpoB* genes were extracted from the trycycler-assembled contigs using the BV-BRC Comparative Systems Service (42). Extracted nucleotide sequences were aligned using Mafft (v7) and visualized on the BV-BRC Multiple Sequence Alignment and SNP / Variation Analysis Service (42,46).

### Phenotype testing of select isolates

The biochemical profile and antimicrobial susceptibility phenotype of a subset of isolates was assessed at the University of Georgia College of Veterinary Medicine Athens Veterinary Diagnostics Laboratory. using the Vitek2 Gram-positive bacteria ID card for biochemical tests and the Sensititre COMPGP1F plates (ThermoFisher) for MIC testing, according to the manufacturers’ instructions.

For MICs, we inoculated sterile H_2_O with *H. ovis* to achieve a 0.5 McFarland; 10 uL of the inoculum was added to 10 mls of Mueller-Hinton broth containing lysed horse blood and supplemented with 0.1 mg of pyridoxal HCL. Finally, 50 uLs of the Mueller-Hinton broth containing *H. ovis* were aliquoted into each well of the Sensititre plate, incubated at 35C in aerobic conditions, and read at 24 and 48 hours.

For biochemical testing on the Vitek2 system, we inoculated 0.45% saline with *H. ovis* to achieve a 0.5 McFarland and entered the cards into the Vitek2 system. The Vitek2 system then made the appropriate dilutions and automatically read them at 15-minute intervals until completed, which was 5 to 8 hours, depending on the isolate. We opted to use the Gram-positive ID card because it contains all of the biochemical tests used to identify *H. ovis* in previous studies (13). For biochemical testing, we selected the 4 *H. ovis* strains with complete genome assemblies (KG36, KG37, KG104, and KG106) and the 4 *Helcococcus* cryptic strains (KG38, KG95, KG105, KG197). For antimicrobial sensitivity testing, we selected 10 *H. ovis* strains representing each tetracycline resistance gene profile including *tetM* only (KG100, KG106, and KG113), *tetA*/*tetB*(KG36, KG92, KG196), *tetT* only (KG104, KG109, KG120), and none (KG107).

### Pangenome Analysis

The *H. ovis* pangenome was constructed using Roary with default parameters and gene clusters were annotated using the BV-BRC Comparative Systems Service (42). To identify gene clusters associated with metritis, we used Scoary with default parameters (20). Scoary identifies gene presence or absence variants significantly associated with a trait by performing Fisher’s Exact Tests. It then uses the phylogenetic relations between strains to look for the causal set of genes. Causal genes were defined as those with Bonferroni-corrected p-values < 0.05.

### Virulome and Resistome

Abricate was used to screen all assembled genomes for ARGs using the NCBI AMRFinder and CARD databases (github.com/tseemann/abricate) (47,48). ARGs associated with point mutations were excluded due to a lack of experimental data for the *Helcococcus* genus. Abricate was also used to screen for virulence factors against the VFDB for known plasmids against the PlasmidFinder database (49,50). Virulence factors were further manually searched for using the BV-BRC Comparative Systems Service (42).

## Data Availability

The whole-genome sequences and the trimmed reads have been uploaded into the NCBI Sequence Read Archive and are found under BioProject number PRJNA514352. SRA accession numbers for the trimmed reads are listed in Supplemental File 2. GenBank accession numbers for the genome assemblies are listed in Supplemental File 3.

## Ethics Declarations

Experimental procedures involving cows were performed in accordance with relevant guidelines and regulations and were approved by the Institutional Animal Care and Use Committee of the University of Florida, under protocol number 201910623. The authors declare no competing interests.

